# Selective degradation of an ER stress-induced protein by ER-associated degradation mechanism during stress recovery

**DOI:** 10.1101/2022.06.06.495033

**Authors:** Yi Wang, Zhihui Ma, Congcong Zhang, Yongwu Chen, Liangguang Lin, Juan Mao, Jianjun Zhang, Linchuan Liu, Pengcheng Wang, Jianming Li

**Affiliations:** University of Chinese Academy of Science, Beijing, 100004, China; Shanghai Center for Plant Stress Biology, The Center of Excellence for Molecular Plant Sciences, Chinese Academy of Sciences, Shanghai, 201602, China; State Key Laboratory for Conservation and Utilization of Subtropical Agro-Bioresources, South China Agriculture University, Guangzhou 510642, China.; Guangdong Key Laboratory for Innovative Development and Utilization of Forest Plant Germplasm, College of Forestry and Landscape Architecture, South China Agricultural University, Guangzhou, Guangdong Province, 510642, China; Department of Molecular, Cellular, and Developmental Biology, University of Michigan, Ann Arbor, MI 48109-1048

**Keywords:** ER-associated degradation, UPR, TIN1, stress recovery, N-glycan

## Abstract

Unfolded protein response (UPR) is a conserved signaling pathway that is activated by accumulation of misfolded proteins in the endoplasmic reticulum (ER) and stimulates production of ER chaperones to restore ER proteostasis. However, little is known how UPR-induced proteins return to their pre-stress levels upon removal of ER stress. TUNICAMYCIN-INDUCED1 (TIN1) is an Arabidopsis protein that is normally expressed in pollen but is rapidly induced by ER stresses in vegetative tissues. Here we show that the ER stress-induced TIN1 is rapidly degraded in the UPR recovery phase. We found that TIN1 degradation depends on its asparagine-linked glycans and requires both EMS-mutagenized bri1 suppressor 5 (EBS5) and EBS6 for its recruitment to the ER-associated degradation (ERAD) complex. Loss-of-function mutations in Arabidopsis ERAD components greatly stabilize TIN1. Interestingly, two other UPR-induced proteins that are coexpressed with TIN1 remained stable upon removal of ER stress, suggesting that rapid degradation during the stress-recovery phase likely applies to a subset of UPR-induced proteins. Further investigation should uncover the mechanisms by which the ERAD machinery differentially recognizes UPR-induced ER proteins.

## Introduction

Endoplasmic reticulum (ER) is an essential eukaryotic cellular organelle that plays an important role in calcium homeostasis, lipid biosynthesis, and folding, modification, complex assembly, and transport of a wide range of transmembrane and secretory proteins (Ferro-Novick et al., 2013). Because protein folding is an error-prone process that can be easily disturbed by various cellular stresses, misfolded proteins often accumulate in the ER lumen, causing the so-called ER stress that interferes with the normal secretory processes crucial for cell survival. As a result, a stress response pathway, widely known as UPR for unfolded protein response, is activated to promote production of protein chaperones, folding catalysts, and components of a unique degradation system known as ER-associated degradation (ERAD) (Hetz et al., 2020). UPR activation leads to translational arrest (reducing the influx of newly synthesized polypeptides into the ER), expansion of the ER membrane and increased production of molecular chaperones and folding catalysts (increasing the protein folding capacity in the ER), and degradation of irreparable misfolded proteins by ERAD, thus re-establishing ER homeostasis and restoring ER function (Hetz et al., 2020). However, little is known how UPR-induced proteins return to their pre-stress levels during the stress recovery phase. Several recent studies suggested involvement of a unique autophagy process, termed as “recov-ER-phagy” in returning the expanded ER to its pre-stress size (Fumagalli et al., 2016; Loi et al., 2019).

The UPR is a highly conserved cellular stress pathway from yeast to plants to human (Hollien, 2013). Yeast is equipped with just one UPR branch consisting of an ER membrane-anchored IRE1 (inositol-requiring enzyme 1), a dual functional Ser/Thr kinase and endoribonuclease that catalyzes a cytosolic splicing reaction to remove an intron of a mRNA to produce its translatable mRNA of HAC1 [homologous to ATF/CREB 1, a yeast basic-leucine zipper (bZIP) protein of 230 amino acids] (Hernandez-Elvira et al., 2018; Adams et al., 2019). Mammalian cells have three UPR branches: IRE1, PERK [protein kinase RNA-like endoplasmic reticulum kinase] and ATF6 (activation transcription factor 6) (Chakrabarti et al., 2011). PERK phosphorylates eIF2α (eukaryotic initiation factor 2α) to globally inhibit protein biosynthesis and reduce the protein load into the ER (Lu et al., 2004), while ATF6 is a membrane-anchored bZIP-type transcription factor that requires two sequential proteolytic cleavages [by site-1 protease (S1P) and site-2 protease (S2P)] on the Golgi membrane to release a transcriptionally-active fragment capable of translocation into the nucleus (Ye et al., 2000). Plants have two UPR branches, IRE1-bZIP60 (a plant equivalent of HAC1) and bZIP28/17 (the Arabidopsis homologs of ATF6), but lack the PERK pathway (Howell, 2013). The yeast and plant UPR systems might rely on IRE1, which can cleave certain mRNAs via regulated-IRE1-dependent decay (RIDD) (Hollien and Weissman, 2006), to reduce global protein synthesis (Mishiba et al., 2013).

In addition to decreasing the protein load and increasing the ER folding capacity, another consequence of UPR activation is enhanced ERAD efficiency. ERAD is a unique degradation mechanism that degrades ER-retained but irreparably misfolded proteins and consists of 4 interdependent steps: recognition/recruitment, ubiquitination, retrotranslocation, and proteasome-mediated proteolysis in the cytosol (Berner et al., 2018). The ERAD machinery is a multi-subunit protein complex that builds around an ER membrane-anchored ubiquitin (E3) ligase. In yeast, at least two ERAD complexes were known: Hrd1 (HMG-reductase degradation1) complex that degrades ERAD-L/M substrates with structural defects in their ER-luminal (L)/membrane(M) domains and Doa10 (Degradation of alpha2 10) complex that degrades ERAD-C clients carrying cytosol (C)-facing structural lesions (Zattas and Hochstrasser, 2015). In Arabidopsis, there are at least three sets of ER-membrane anchored E3 ligases: AtHrd1a/b (two homologs of the yeast Hrd1), Doa10a/b (two homologous to the yeast Doa10), and RMA1-3 (Ring membrane-anchored1-3) (Chen et al., 2020). The Arabidopsis Hrd1-containing ERAD complex contains several conserved proteins, AtHrd1a/b (Su et al., 2011), EBS6 (EMS-mutagenized bri1 suppressor6)/AtOS9 (*Arabidopsis thaliana* homolog of the mammalian osteosarcoma amplified-9) (Huttner et al., 2012; Su et al., 2012), EBS5/AtSel1A (*Arabidopsis thaliana* homolog of the mammalian Sel1 for Suppressor of lin-12-like 1)(Liu et al., 2011; Su et al., 2011), an ER membrane-anchored ubiquitin conjugase UBC32 (U-box containing protein32) (Cui et al., 2012), and two plant-specific components, EBS7 and PAWH1/2 (Protein Associated With Hrd1 1/2), which are important for regulating the stability of AtHrd1 (Liu et al., 2015; Lin et al., 2019).

TUNICAMYCIN-INDUCED 1 is an Arabidopsis protein that is rapidly induced by several chemical inducers of ER stress (Iwata et al., 2010), such as tunicamycin (TM, a widely used inhibitor of the biosynthesis of the lipid-linked N-glycan precursor (Duksin and Mahoney, 1982), dithiothreitol (DTT, a strong reducing agent capable of disrupting disulfide bridges of newly synthesized and folded proteins), and azetidine-2-carboxylate [AZC, a proline analog whose incorporation into proteins causes protein misfolding (Trotter et al., 2001)]. TIN1 is a plant-specific protein that lacks any known motif or domain and is normally expressed in pollen (Iwata et al., 2012). Previous studies with loss-of-function and overexpression approaches showed that TIN1 is involved in the formation of pollen surface structure (Iwata et al., 2012; Iwata et al., 2017). However, it remains unknown about its biochemical function and its post-stress recovery mechanism. Given its rapid induction by ER stress inducers and its lack of known domain or motif involved in protein folding and quality control, it was previously hypothesized that TIN1 might function as a protein cochaperone to assist protein folding, repair, and/or degradation (Iwata et al., 2010). In this study, we initially discovered TIN1 as a potential Hrd1-interacting ERAD component in Arabidopsis. However, our genetic and biochemical experiments showed that TIN1 is not an ERAD component but rather an ERAD substrate, which is rapidly degraded via the Hrd1-containing ERAD complex in an asparagine-linked glycan (N-glycan) dependent manner. Interestingly, we found that neither BIP3 (binding immunoglobulin protein 3) nor AtERdj3A (*Arabidopsis thaliana* ER-localized DnaJ-domain protein 3A), two other well-known UPR-induced proteins, remain very stable upon removal of ER stress. Further investigation is needed to understand the biochemical basis for their differential post-ER-stress fates.

## Results

### Identification of TIN1 as an Hrd1-associated protein by immunoprecipitation-mass spectrometry

A previous immunoprecipitation-mass spectrometry experiment, which identified PAWH1/2 as plant-specific components of the Arabidopsis ERAD complex (Lin et al., 2019), discovered TIN1 as a Hrd1-associated protein. As its name indicates, TIN1 was originally identified as an Arabidopsis protein that is highly induced by TM (Iwata et al., 2010). Consistent with earlier studies, the *TIN1* transcript was highly induced by several ER stress-inducers including TM, DTT, and AZC, which were revealed by a quantitative real-time reverse transcription PCR (qRT-PCR) analysis and histochemical staining of young seedlings of a transgenic Arabidopsis line expressing a *pTIN1::GUS* reporter construct consisting of a *TIN1* promoter and the cDNA of the reporter β-glucuronidase (**Supplemental Figure S1, A and B**). Our qRT-PCR analyses also revealed that the *TIN1* transcript was induced by the plant stress hormone abscisic acid (ABA), mannitol (used widely as an osmotic stress inducer), and the 37°C heat treatment (**Supplemental Figure S1, C-E**). Interestingly, the salt treatment, which was known to activate an important UPR regulator bZIP17 (Liu et al., 2007), had little impact on the *TIN1* transcript abundance (**Supplemental Figure S1F**). We also analyzed the *TIN1* transcript abundance in TM-treated seedlings of the wild-type Arabidopsis and several previously-characterized UPR mutants and found that the ER stress-induced *TIN1* expression is likely mediated by the two known branches of the Arabidopsis UPR mechanism as the TM-induction of the *TIN1* transcript was reduced in the 4 analyzed Arabidopsis UPR mutants, including *ire1a ire1b* lacking the two Arabidopsis homologs of the yeast IRE1, *s2p* missing the Golgi-localized site-2 protease involved in cleaving and activating bZIP17 and bZIP28, *bzip28*, and *bzip60* mutants (**Supplemental Figure S1G**).

To study the impact of ER stress on the TIN1 protein level, we generated an anti-TIN1 antibody that detected the presence of TIN1 protein in DTT- or TM-treated wild-type seedlings and a TIN1-GFP fusion protein in a *p35S::TIN1-GFP* transgenic Arabidopsis line but not in stressed seedlings of a previously characterized T-DNA insertional *tin1-1* mutant (Iwata et al., 2012) (**Supplemental Figure S2, A and B**). As shown in Figure 1**, A-C**, treatment of DTT, TM, or AZC resulted in significant accumulation of the TIN1 protein. It is worth noting that the molecular weight of the TM-induced TIN1 protein is smaller than that of the DTT/AZC-induced TIN1 protein, likely due to the TM-induced inhibition of N-glycosylation. Consistent with the qRT-PCR results and our *pTIN1::GUS* histochemical staining (**Supplemental Figure S2C**), our immunoblot analysis revealed that TIN1 can only be detected in the pollen of the Arabidopsis plants grown under normal growth conditions (**Supplemental Figure S2D**).

**Figure 1.**
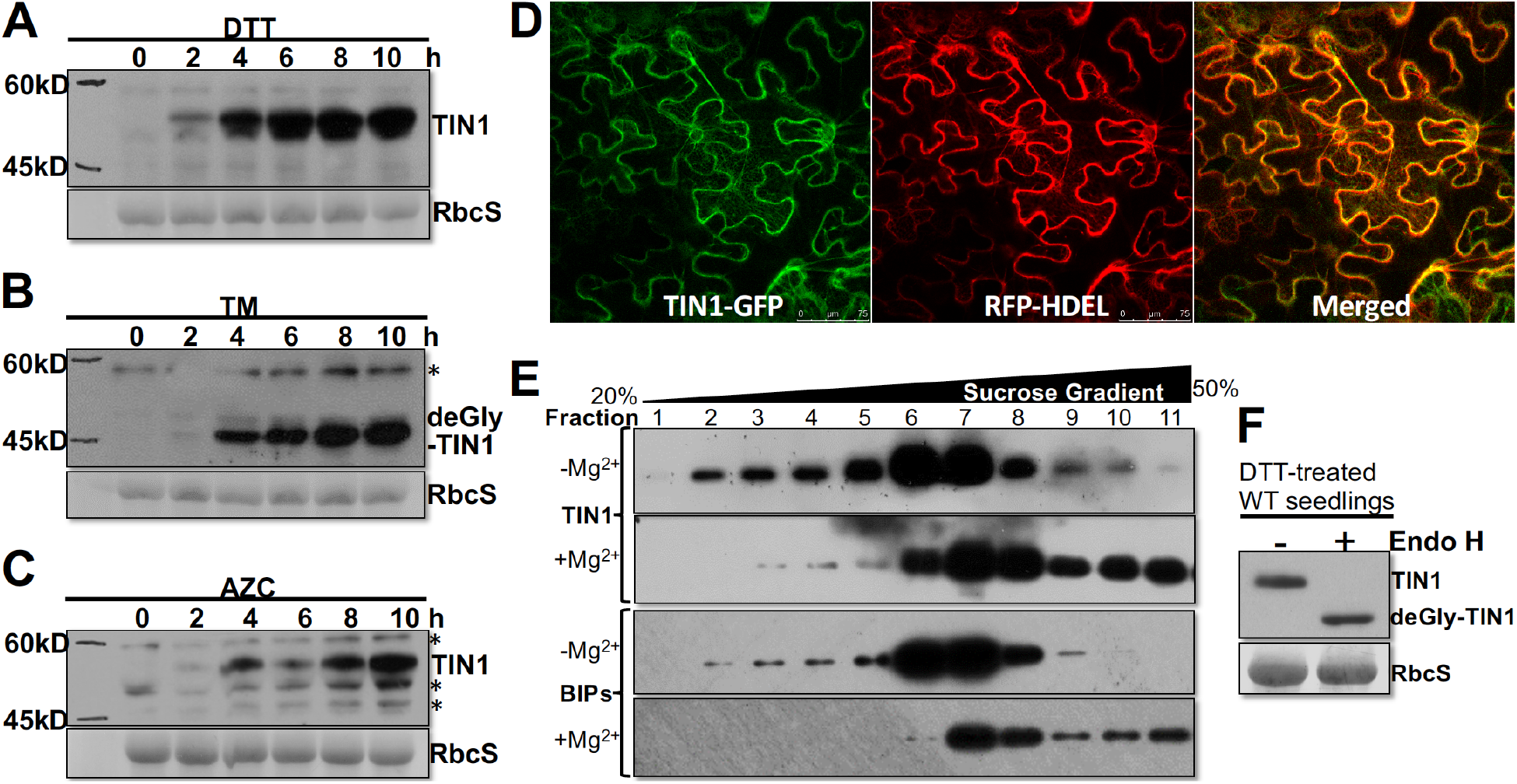
The TIN1 protein is an ER-localized protein that can be induced by ER stress. **A-C**. Immunoblot analysis of the TIN1 accumulation in Arabidopsis seedlings treated with TM (**A**), DTT (**B**), and AZC (**C**). **D**. Confocal analysis of the subcellular localization of a transiently expressed TIN1-GFP fusion protein in tobacco leaf epidermal cells. **E**. A sucrose gradient ultracentrifugation analysis of the endogenous TIN1 protein. **F**. The Endo H analysis of the endogenous TIN1 protein. The wild-type seedlings used in **E** and **F** were pretreated with DTT for 4 hours to increase the TIN1 abundance. In **A**-**C** and **F**, the ponceau red-stained RbcS serves as the sample loading control.

In agreement with an earlier report (Iwata et al., 2010), TIN1 is an ER-localized protein despite its lack of the HDEL (His-Asp-Glu-Leu) ER retrieval motif at its C-terminus. As shown in Figure 1D, a TIN1-GFP (green fluorescent protein) fusion protein exhibited an overlapping fluorescent pattern with HDEL-tagged red fluorescent protein (RFP-HDEL), a widely used ER marker protein (Liu and Dixon, 2001), when they were coexpressed in tobacco (*Nicotiana benthamiana*) leaf epidermal cells. To eliminate the possibility that the overlapping ER localization patterns of TIN1-GFP and RFP-HDEL could be an artifact due to overexpression of a GFP-tagged fusion protein that carries a predicted N-terminal signal peptide in tobacco leaf epidermal cells (Figure 1D) or in Arabidopsis protoplasts (Iwata et al., 2010), we performed a sucrose gradient ultracentrifugation experiment in the presence or absence of Mg^2+^, which was known to be required for ribosome-ER association (Lord et al., 1973), with protein extracts of DTT-treated wild-type Arabidopsis seedlings. The DTT treatment was necessary to detect the TIN1 protein in Arabidopsis vegetative tissues. As shown in Figure 1E, addition of Mg^2+^ shifted the elution profile of the endogenous TIN1 protein towards higher density, similar to that of BIP1, a known ER-localized heat shock protein 70 (HSP70). Additional support for its ER localization was obtained by a simple biochemical assay utilizing endoglycosidase H (Endo H), which removes high mannose-type N-glycans of ER localized or retained glycoproteins but not Golgi-processed complex-type N-glycans of glycoproteins that traffic through the Golgi apparatus (Tarentino et al., 1974). As shown in Figure 1F, the endogenous TIN1 protein of DTT-treated wild-type Arabidopsis seedlings was sensitive to Endo H. Taken together, our experiments demonstrated that the endogenous TIN1 protein is an ER-localized glycoprotein.

### Neither TIN1 nor its likely homolog is involved in ERAD

Because of its coimmunoprecipitation with Hrd1 and its ER localization, we initially hypothesized that TIN1 might be an ER luminal component of the Hrd1-containing ERAD complex, which was shown to be involved in the degradation of two ER-retained mutant variants of the brassinosteroid (BR) receptor BRASSINOSTEROID-INSENSITIVE1 (BRI1), including bri1-5 and bri1-9 (Chen et al., 2020). Our previous studies showed that mutations in components of the Hrd1-containing ERAD complex greatly stabilized the two mutant bri1 receptors and partially rescued the growth phenotypes of the corresponding dwarf mutants due to leakage of ER-accumulated bri1-5 or bri1-9 proteins to the plasma membrane (PM) (Su et al., 2011, 2012; Liu et al., 2015; Lin et al., 2019). If TIN1 were a true component of the Arabidopsis Hrd1 complex, a loss-of-function *tin1* mutation would inhibit degradation of the two mutant proteins and suppress the dwarf phenotypes of *bri1-5* and *bri1-9*. To test this hypothesis, we obtained a previously characterized T-DNA insertional mutant of TIN1, *tin1-1* (Iwata et al., 2012) that lacks a detectable amount of TIN1 protein (**Supplemental Figure S2B**), and crossed it with *bri1-5* and *bri1-9* to generate the *tin1-1 bri1-5* and *tin1-1 bri1-9* double mutants. We found that *tin1-1* had little effect on the growth morphology of the *bri1-5* or *bri1-9* mutant and failed to increase the abundance of the mutant bri1-5 and bri1-9 proteins (Figure 2, A **and** B). It is important to note that despite repeated electron microscopic observations, the *tin1-1* mutant or the *TIN1-GFP* overexpression line failed to exhibit any observable difference in the surface of pollen grains compared to the wild-type Arabidopsis plants (**Supplemental Figure S3**), a phenotype that was previously observed in *tin1-1* mutants or *TIN1*-overexpressing transgenic Arabidopsis lines (Iwata et al., 2012; Iwata et al., 2017). We suspected that our failure to detect the pollen wall phenotype might be caused by our growth conditions that were different from that used in the two previously published studies.

**Figure 2.**
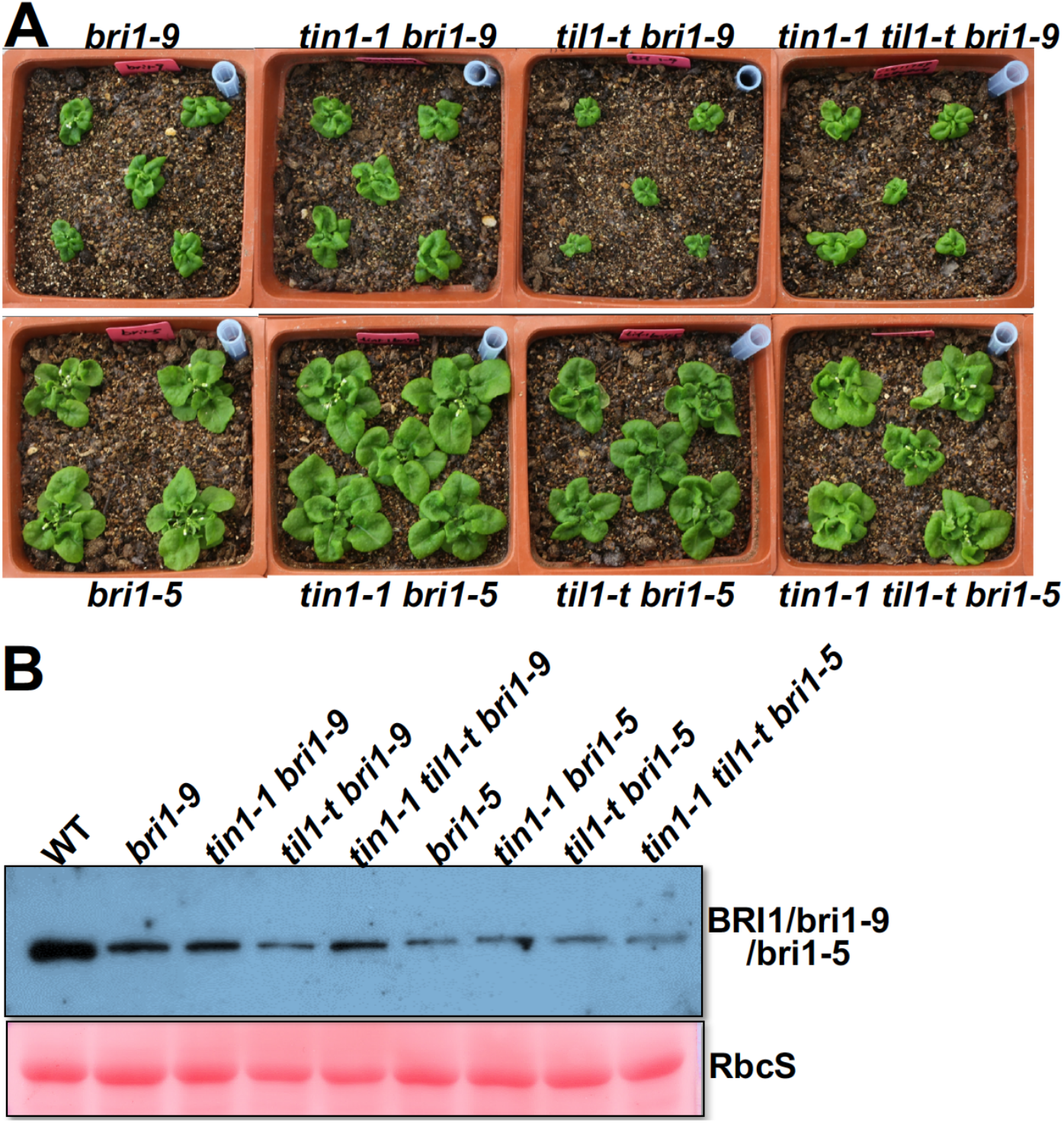
Neither TIN1 nor TIL1 is involved in ERAD of bri1-5 or bri1-9. **A**. Pictures of 2-week-old soil-grown plants of *bri1-5*, *bri1-9*, and various double and triple mutants. **B**. Immunoblot analysis of BRI1 abundance in 2-week-old seedlings of various genotypes shown in **A.** The ponceau red-stained RbcS serves as the loading control.

We thought that the failure of the *tin1-1* mutation to inhibit degradation of bri1-5 and bri1-9 might simply be caused by functional redundancy between TIN1 and a potential TIN1-like protein (At1g47310, named hereinafter TIL1 for TIN1-Like1). TIL1 is a 395-amino acid polypeptide exhibiting 22%/58% sequence similarity/identity with TIN1 (**Supplemental Figure S4A**), and the presence of the TIN1-TIL1 pair is widely detected in the sequenced genomes of seed plants (**Supplemental Figure S4B**). Similar to TIN1, TIL1 is likely localized in the ER as revealed by confocal microscopy of a TIL1-GFP fusion protein transiently expressed in tobacco leaves (**Supplemental Figure S5A**) and sucrose-gradient ultracentrifugation analyses of the TIL1-GFP fusion protein that was expressed in transgenic Arabidopsis plants (**Supplemental Figure S5B**). It is important to note that *TIL1* is a constitutively expressed gene and is insensitive to TM treatment (**Supplemental Figure S5, C and D**). Consistent with its ER localization, the TIL1-GFP fusion protein is sensitive to Endo H digestion (**Supplemental Figure S5E**).

To test the potential functional redundancy of TIN1 and TIL1 in an Arabidopsis ERAD process, we obtained a T-DNA insertional mutant *til1-t* from ABRC (*SAIL_165_E10* containing a T-DNA insertion in the 1st exon),which is morphologically indistinguishable from the *tin1-1* mutant or its wild-type control (**Supplemental Figure S6**), and generated *til1-t bri1-9* and *tin1-1 til1-t bri1-9* mutants. Unfortunately, neither the *til1-t* nor the *tin1-1 til1-t* double mutation had a detectable effect on the growth morphology of the *bri1-5* and *bri1-9* mutants or the protein abundance of the corresponding mutant bri1 proteins (Figure 2, A **and** B). It is worthy to note that the tin1-1 til1-t double mutant also failed to display the pollen surface defect under our experimental conditions (**Supplementary Figure 3**). Consistent with the mutant analyses, overexpression of TIN1 or TIL1 in *bri1-9* had no obvious impact on the growth phenotype of the dwarf mutant (**Supplemental Figure S7, A and B**). These data argued strongly against the hypothesis that TIN1 and TIL1 are functionally redundant components of the Arabidopsis Hrd1-containing ERAD complex that degrades the ER-retained mutant BR receptors. It is important to point out that our results did not completely eliminate a possibility that TIN1 and TIL1 might be involved in an ERAD process that degrades yet to be discovered ERAD clients in Arabidopsis. A quantitative proteomic study using total proteins of the *tin1-1 til1-t* double mutant and its wild-type control could help to investigate such a possibility.

### TIN1 is rapidly degraded while TIL is a stable protein

Our failure to demonstrate involvement of TIN1 and/or TIL1 in an ERAD process prompted us to investigate an alternative hypothesis of TIN1 being a potential ERAD client. To test this, we performed a cycloheximide (CHX)-chasing experiment to analyze the stability of the DTT-induced TIN1 protein as CHX is a widely used protein biosynthesis inhibitor. Because TIN1 is barely detectable in vegetative tissues under normal growth conditions (**Supplemental Figure S2**), a 4-h pretreatment with DTT is necessary to increase its transcription to permit easy detection of its protein product for the CHX-chasing experiment (**Supplemental Figure S2A**). As shown in Figure 3A, The DTT-induced TIN1 is rapidly degraded with its half-life being ∼30 min. Additional support for TIN1 being a very unstable protein came from our analysis of the TIN1 abundance of Arabidopsis seedlings treated with DMSO or MG132, a widely used inhibitor of proteasome. As shown in Figure 3B, the DMSO-treated seedlings accumulated no detectable amount of TIN1 protein, whereas the MG132 treatment resulted in great accumulation of TIN1. We also tested the protein stability of AZC-induced TIN1. As shown in **Supplemental Figure S8**, the AZC-induced TIN1 protein was also rapidly degraded upon AZC removal. Because both DTT and AZC can cause TIN1 misfolding or disrupt TIN1 interaction with other ER-localized proteins to reduce its protein stability, we also analyzed the stability of a TIN1-GFP fusion protein in a *p35S::TIN1-GFP* transgenic line using the CHX-chasing approach and found that TIN1-GFP was also an unstable protein with a half-life of ∼ 3 hours (Figure 3C). Similarly, treatment with MG132 also increased the stability of the TIN1-GFP fusion protein in the absence or presence of CHX (Figure 3, D **and** E). It is interesting to note that the stability of TIN1-GFP is stronger than that of the DTT/AZC-induced endogenous TIN1, likely caused by the strong *p35S* promoter or the C-terminal GFP tag that might interfere with TIN1 degradation. Taken together, these results demonstrated that TIN1 is a highly unstable protein that is likely degraded by the cytosolic proteasome in vegetative tissues. By contrast, TIL1 is a very stable protein. A CHX-chasing experiment revealed a little change in the abundance of TIL1-GFP in *p35S::TIL1-GFP* transgenic seedlings (**Supplemental Figure S9**).

**Figure 3.**
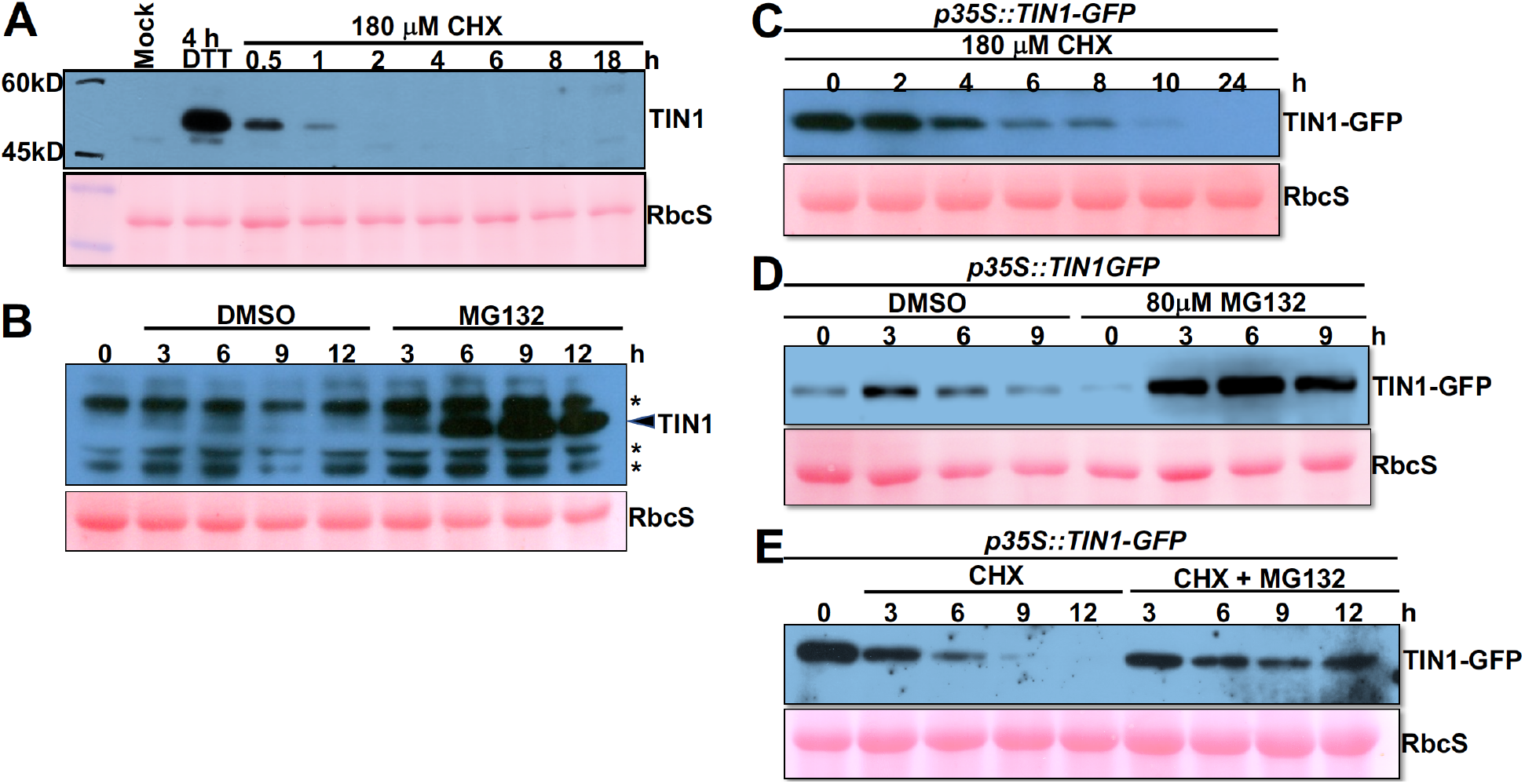
TIN1 is rapidly degraded via a proteasome-mediated process. **A**. Immunoblot analysis of the TIN1 stability in seedlings treated with DTT (for 4 hours) followed by CHX for varying duration. **B**. Immunoblot analysis of the TIN1 protein abundance in seedlings treated with DMSO or MG132. **C**. Immunoblot analysis of the protein stability of a TIN1-GFP fusion proteins in *p35S::TIN1-GFP* transgenic seedlings treated with 180 μM CHX for different durations. **D-E**. Immunoblot analysis of the TIN1-GFP protein abundance in *p35S::TIN1-GFP* seedlings treated with DMSO or MG132 in the presence or absence of CHX.

### TIN1 degradation relies on its N-glycans

Because TIN1 is an N-glycosylated protein (Figure 1F), we investigated whether its rapid degradation upon removal of an ER stressor depends on its N-glycan. We first treated the wild-type Arabidopsis seedlings with DTT to induce TIN1 accumulation, followed by treatment with CHX in the presence or absence of kifunensine (Kif), a well-known inhibitor of ERAD by blocking α1,2-mannosidases that generate the conserved N-glycan ERAD signal (Elbein et al., 1990). As shown in Figure 4, A **and** B, TIN1 remained detectable 9 h after CHX addition in the presence of Kif, whereas the TIN1 abundance was markedly reduced 1 h after CHX addition in the absence of Kif, suggesting that TIN1 degradation is likely N-glycan-dependent. A similar experiment performed with seedlings of the *p35S::TIN1-GFP* transgenic line also revealed that Kif effectively prevented the CHX-triggered disappearance of TIN1-GFP (Figure 4C).

**Figure 4.**
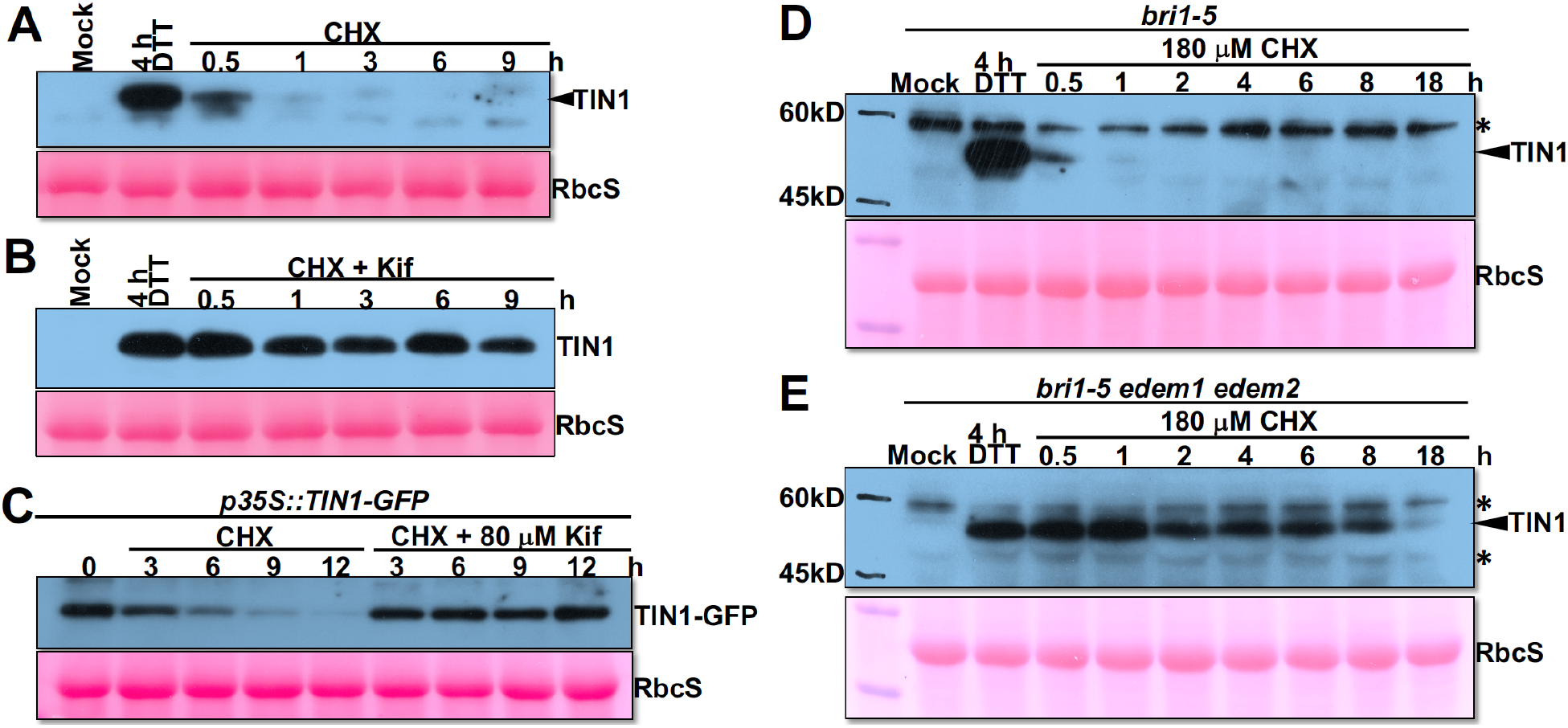
TIN1 degradation relies on its N-glycans. **A-B.** Immunoblot analysis of the TIN1 protein abundance in 2-week-old seedlings treated with DTT (for 4 hours) followed by CHX (for varying durations) in the absence (**A**) or presence (**B**) of Kif. **C**. Immunoblot analysis of the TIN1-GFP abundance in 2-week-old *p35S::TIN1-GFP* transgenic seedlings treated with CHX for different durations in the presence or absence of Kif. **D-E**. Immunoblot analysis of the TIN1 protein level in 2-week old seedlings of *bri1-5* (**D**) or *bri1-5 edem1 edem2* triple mutant (**E**), which were treated with DTT for 4 hours followed by treatment with 180 μM CHX for varying durations.

A further support for the N-glycan-dependent degradation of TIN1 came from our analysis of the TIN1 stability in an Arabidopsis double mutant lacking the two Arabidopsis homologs of the mammalian EDEMs, AtEDEM1 and AtEDEM2 (known previously as MNS4 and MNS5 for mannosidase 5 and 4 (Huttner et al., 2014), respectively]. An earlier study showed that AtEDEM1/MNS5 and AtEDEM2/MNS4 function redundantly in degrading the ER-retained bri1-5 (Huttner et al., 2014). As shown in Figure 4, D **and** E, a CHX-chasing experiment with *bri1-5* mutant seedlings showed a rapid degradation of the DTT-induced TIN1, whereas a similar CHX-chasing experiment performed with seedlings of the *edem1 edem2 bri1-5* triple mutant showed that the DTT-induced TIN1 protein remained stable for at least 8 hours after the CHX treatment. Consistent with the inhibitory impact of the *edem1 edem2* double mutation on TIN1 degradation, TIN1 was found to interact with AtEDEM2/MNS4 when the two proteins were coexpressed as fusion proteins in leaf epidermal cells of tobacco (*Nicotiana benthamiana*), as revealed by a bimolecular fluorescence complementation (BiFC) assay and a coimmunoprecipitation (coIP) experiment (**Supplemental Figure S10, A and B**). These results allowed us to conclude that TIN1 degradation is largely dependent on its N-glycans.

### Both N^296^ and N^406^ are important for TIN1 degradation

Sequence analysis indicated that TIN1 has three potential N-glycosylation sites: N^197^ (N for asparagine), N^296^, and N^406^ (**Supplemental Figure S4A**). To determine whether or not all three predicted N-glycan sites are indeed glycosylated and involved in TIN1 degradation, we generated Arabidopsis transgenic lines expressing one of the seven TIN1-GFP fusion proteins lacking 1 (N197Q, N296Q, N406Q; Q for glutamine), 2 (N197,296Q, N197,406Q, N296,406Q), or all 3 [N197,296,406Q (N3Q)] potential N-glycosylation sites, and subsequently analyzed the impact of these 7 mutations on the mobility and Endo H or Kif sensitivity of mutant TIN1-GFP fusion proteins. Representative transgenic lines were used for the biochemical assays. As shown in Figure 5A, all 7 mutations resulted in faster mobility of the TIN1-GFP fusion proteins on SDS-PAGE. It is interesting to note that the three single N-Q mutant TIN1-GFP fusion proteins exhibited different mobility on the SDS-PAGE gel with TIN1^N406Q^-GFP running a bit slower than TIN1^N197Q^-GFP and TIN1^N296Q^-GFP, which is likely caused by its position near the C-terminus. Consistent with this observation, the TIN1^N197,406Q^-GFP and TIN1^N296,406Q^-GFP bands moved a bit slower than the TIN1^N197,296Q^-GFP band on the SDS-PAGE gel. In line with the mobility difference of the 7 mutant TIN1-GFP fusion proteins, the Endo H treatment showed that while the Endo H digestion caused no change in the mobility of the TIN1^N3Q^-GFP, it did result in mobility change of the three double N-Q mutant TIN1-GFP fusion proteins on SDS-PAGE gel (Figure 5B). These experiments confirmed that TIN1 is N-glycosylated at the three predicted N residues.

**Figure 5.**
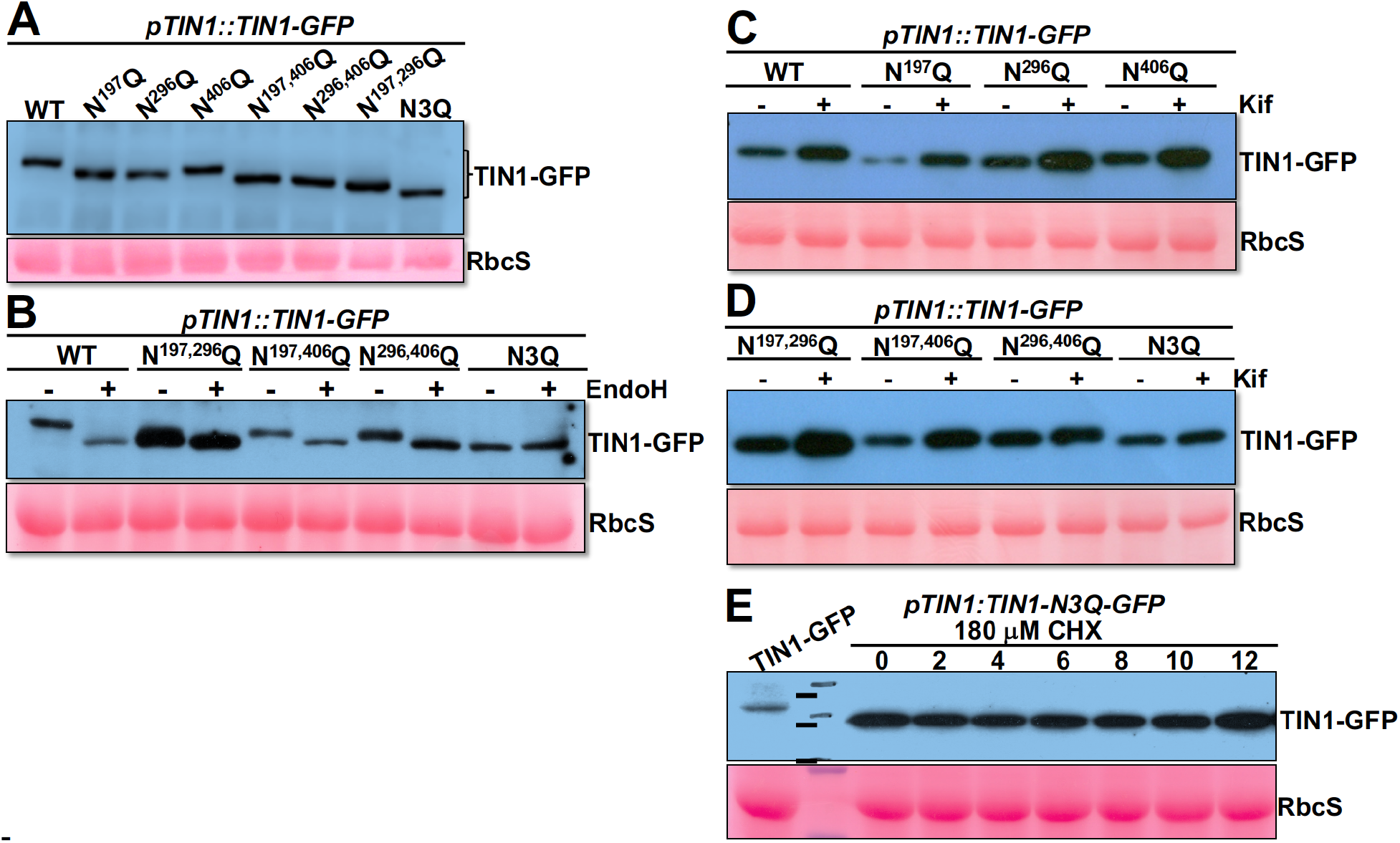
Both Asn^296^ and Asn^406^ are important for the rapid degradation of TIN1. **A**. Analysis of the Asn-Ala mutations on the electromobility of the TIN1-GFP protein on SDS-PAGE. **B**. The Endo H sensitivity of various Asn-Ala mutated TIN1-GFP proteins. **C-D**. The Kif sensitivity of various Asn-Ala-mutated TIN1-GFP proteins. **E**. The immunoblot analysis pf the TIN1-GFP(N3Q) protein abundance in two week-old *p35S::TIN1-GFP(N3Q)* seedlings treated with 180 μM CHX for varying durations. In **A**-**E**, the ponceau red-stained RbcS serves the loading control.

We also treated those transgenic seedlings with 25 μM Kif for 6 hours and analyzed the TIN1 protein abundance by immunoblotting. As shown in Figure 5C, the Kif treatment seemed to have a stronger impact on TIN1^N197Q^-GFP than TIN1^N296Q^-GFP or TIN1^N406Q^-GFP. Consistently, Kif increased the protein abundance of TIN1^N197,296Q^-GFP and TIN1^N197,406Q^-GFP proteins but had a much weaker effect on the protein abundance of TIN1^N296,406Q^-GFP and TIN1^N3Q^-GFP (Figure 5D). These results suggest that N^296^ and N^406^ have a stronger impact on TIN1 degradation than N^197^. More importantly, a CHX-chasing experiment with seedlings of the *p35S:TIN1^N3Q^-GFP* transgenic line indicated that simultaneous elimination of all three N-glycosylation sites effectively blocked the TIN1 degradation (Figure 5E). These results not only confirmed the N-glycan dependence of TIN1 degradation but also revealed potential functional difference among the three glycosylated N residues. It is interesting to note that our sequence analysis showed that the three predicted N-glycosylation sites are highly conserved among the plant TIN1 homologs with the N^197^ residue being the least conserved and only present in TIN1 homologs of angiosperms (Iwata et al., 2010) (**Supplemental Figure S11**).

### TIN1 degradation requires both EBS6 and EBS5

Our results of the Kif treatment, the Arabidopsis *edem1 edem2 bri1-5* mutant, the tobacco TIN1-AtEDEM2/MNS4 interaction experiments, and the Arabidopsis transgenic lines expressing loss-of-glycosylation mutant TIN1-GFP fusion proteins strongly suggested that TIN1 degradation likely requires an ERAD N-glycan signal with an exposed α1,6-mannose residue (Hong et al., 2009; Hong et al., 2012). Previous studies showed that such an ERAD signal is recognized in Arabidopsis by a highly conserved Arabidopsis protein EBS6 (EMS-mutagenized suppressor6) (Su et al., 2012), which is also known as AtOS9 for its sequence similarity to the mammalian OS9 and the yeast yOS9 (yeast OS9 homolog) (Huttner et al., 2012). We thus predicted that EBS6 is required for the TIN1 degradation. To test our prediction, we performed a CHX-chasing experiment with seedlings of a previously described *ebs6-2* mutant and its corresponding wild-type (Su et al., 2012) and found that the *ebs6-2* mutation greatly stabilized the DTT-induced TIN1 protein (Figure 6B) compared to that of the wild-type seedlings (Figure 6A). It is interesting to note that the *ebs6-2* mutation did not completely block the TIN1 degradation as TIN1 was not detectable in *ebs6-2* seedlings 18 h after the CHX treatment (Figure 6B). This finding seemed to be consistent with the result of the CHX-chasing experiment of the *edem1 edem2* double mutant that only had residual amount of TIN1 protein 18 h after the CHX addition (Figure 4E). Our finding also agrees with earlier yeast studies showing that misfolded glycoproteins could still be degraded, albeit at a greatly reduced rate, in the yeast *Δyos9* mutant (Szathmary et al., 2005).

**Figure 6.**
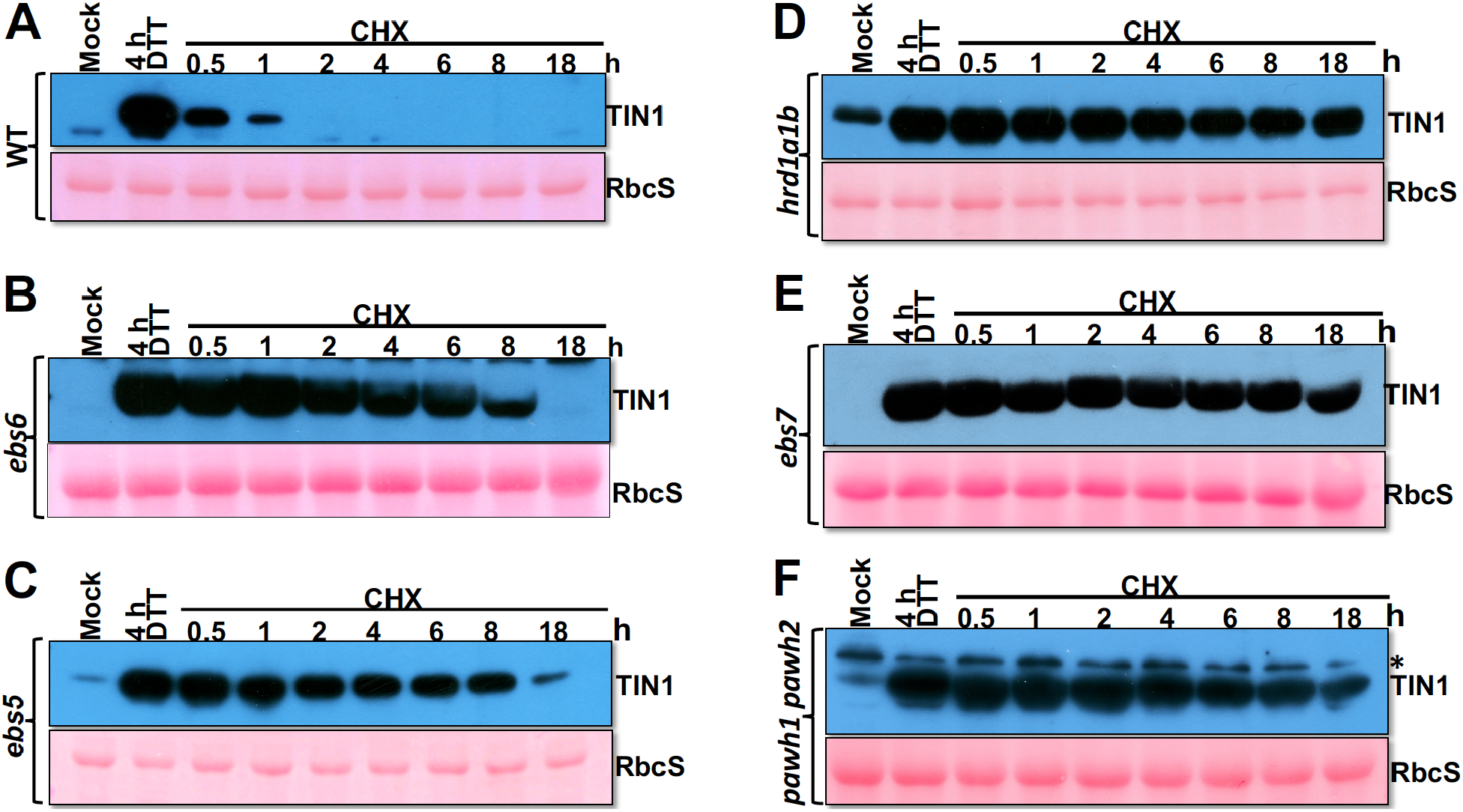
TIN1 degradation requires components of the Arabidopsis Hrd1 complex. **A-F**. Immunoblot analysis of the TIN1 protein abundance in 10-day-old seedlings of indicated genotypes, including wild-type, *ebs5*, *ebs6*, *hrd1a1b*, *ebs7*, and *pawh1 pawh2*, which were treated with DTT (for 4 hours) followed by treatment with 180 μM CHX for varying durations. The ponceau red-stained RbcS serves as the loading control in each immunoblot assay.

A similar CHX study was also performed with seedlings of another Arabidopsis ERAD mutant, *ebs5*, which carries a T-DNA insertional mutation of EBS5 (Su et al., 2011), the Arabidopsis homolog of the yeast Hrd3/mammalian Sel1L that are known to interact with yOS9/OS9 to recruit a terminally misfolded glycoprotein to the ER membrane-anchored E3 ligase for retrotranslocation and ubiquitination (Liu et al., 2011; Su et al., 2011). Similar to what was found in the *ebs6-2* mutant, the stability of the DTT-induced TIN1 was greatly stabilized in the *ebs5-3* mutant (Figure 6C). Together, these biochemical experiments demonstrated that degradation of the DTT-induced TIN1 requires the two interacting recruitment factors known to bring a committed glycosylated ERAD client to the ER membrane-anchored ERAD machinery.

### TIN1 degradation is mediated by the EBS7-PAWH-Hrd1 complex

Because TIN1 is a luminal HDEL-lacking glycoprotein that requires EBS5 and EBS6 for its degradation, we suspected that TIN1 likely uses the ER membrane-anchored E3 ligase Hrd1 complex for its retrotranslocation and ubiquitination. Our previously studies demonstrated the redundant function of the two Arabidopsis homologs of the yeast Hrd1, AtHrd1a and AtHrd1b, in ERAD of bri1-5 and bri1-9 (Su et al., 2011). To investigate a potential role of AtHrd1a/1b in TIN1 degradation, we first examined the TIN1 abundance in several known ERAD mutants, including an Arabidopsis *hrd1a hrd1b* double mutant, and found that TIN1 was easily detected in the *hrd1a hrd1b* double mutant that was not treated with an ER stress-inducing chemical (**Supplemental Figure S12A**). We believed that this was likely caused by the *hrd1a hrd1b*-blocked degradation of the TIN1 protein rather by the *hrd1a hrd1b*-triggered increase in *TIN1* transcript (**Supplemental Figure S12B**). By comparison, the heat treatment caused a much greater increase [∼40 fold (**Supplemental Figure S1E**)] in the *TIN1* transcript abundance than the *hrd1a hrd1b* double mutation [∼4 fold (**Supplemental Figure S12B**)], yet the heat-treated seedlings failed to accumulate detectable amount of TIN1 protein (**Supplemental Figure S13A**). A further support for our conclusion was provided by the CHX-chasing experiments that revealed a very stable TIN1 protein in CHX-treated or heat-stressed *hrd1a hrd1b* mutant seedlings (Figure 6D**; Supplemental Figure S13B**) and a very stable TIN1-GFP fusion protein in *p35S::TIN1-GFP hrd1a hrd1b* transgenic line (**Supplemental Figure S13C**).

Our recent studies discovered that the protein stability and/or activity of the Arabidopsis Hrd1 is regulated by two plant-specific components of the Arabidopsis Hrd1-containing ERAD machinery, EBS7 and PAWH1/2 (Liu et al., 2015; Lin et al., 2019). Consistently, our immunoblot analysis of the TIN1 abundance in *ebs7* and *pawh1 pawh2* mutants showed increased TIN1 abundance in the two ERAD mutants (**Supplemental Figure S12A**). Interestingly, the *ebs6* mutant accumulated less amount of TIN1 protein compared to other ERAD mutants, which is likely caused a potential TIN1-Hrd1 direct interaction. The increased TIN1 abundance in *ebs5* mutant at 18 hour after CHX treatment in comparison to the *ebs6* mutant (Figure 6, B **and** C) could be caused by a combination of no EBS5 and significantly reduced EBS6 level in *ebs5* mutant (Su et al., 2012). We have previously shown that neither *ebs5* nor *ebs6* mutation had an obvious impact on the protein abundance of EBS7, PAWH1/2, or Hrd1 (Lin et al., 2019). We also performed the CHX-chasing experiment with seedlings of *ebs7*-*1* and *pawh1 pawh2* mutants that were known to be defective in ERAD of two mutant bri1 proteins (Liu et al., 2015; Lin et al., 2019). As shown in Figure 6, E **and** F, both *ebs7* and *pawh1 pawh2* mutations greatly stabilized the DTT-induced TIN1 protein. Similar CHX-chasing experiments were conducted with AZC-treated seedlings of *ebs7* and *hrd1a hrd1b* mutants, which revealed that both the *ebs7* and *hrd1a hrd1b* mutation almost completely blocked degradation of the AZC-induced TIN1 (**Supplemental Figure S14, A and B**). Taken together, these biochemical analyses indicated that the ER-stress induced TIN1 is degraded by the EBS7-PAWH1/2-Hrd1 ERAD machinery.

### BIP3 and TMS1 are highly stable proteins

Our finding that an UPR-induced protein is rapidly degraded via ERAD prompted us to examine the stability of two other UPR-induced Arabidopsis proteins upon resolution of the ER stress. Previous studies showed that *BIP3* and *TMS1* (Thermosensitive Male Sterile1) were rapidly and greatly induced by several ER stress inducers (Noh et al., 2003; Iwata and Koizumi, 2005; Yamamoto et al., 2008; Yang et al., 2009; Iwata et al., 2010) and that these two genes were known to be coexpressed with *TIN1* (**Supplemental Figure S15**). BIP3 is one of the three ER-localized HSP70s in Arabidopsis (Noh et al., 2003), while TMS1 is a member of the ER-localized HSP40s functioning as co-chaperones of HSP70s (Yamamoto et al., 2008). TMS1 is also known as AtERdj3A because it is one of the two Arabidopsis homologs of the mammalian ERdj3 (ER-localized DnaJ-like protein3) (Shen and Hendershot, 2005), and was recently shown to interact with BIP3 (Ma et al., 2015). In line with previous results, the transcript abundance and protein levels of BIP3 and TMS1/AtERdj3A were greatly upregulated by DTT and TM (Figure 7, A-C). To investigate if BIP3 and TMS1/AtERdj3A were also rapidly degraded upon removal of DTT, we performed similar DTT-CHX time-course analyses for BIP3 and AtERdj3A with custom-made antibodies. As shown in Figure 7, D **and** E, both BIP3 and TMS1/AtERDj3A remained very stable 24 h after CHX treatment of the DTT-pretreated seedlings. Thus, the rapid degradation via ERAD upon removal of ER stress is not a common recovery mechanism for UPR-induced proteins but likely a unique property of TIN1 or only applies to a subset of UPR-induced proteins, raising an interesting question on what unique feature makes TIN1 a target of the Arabidopsis ERAD machinery.

**Figure 7.**
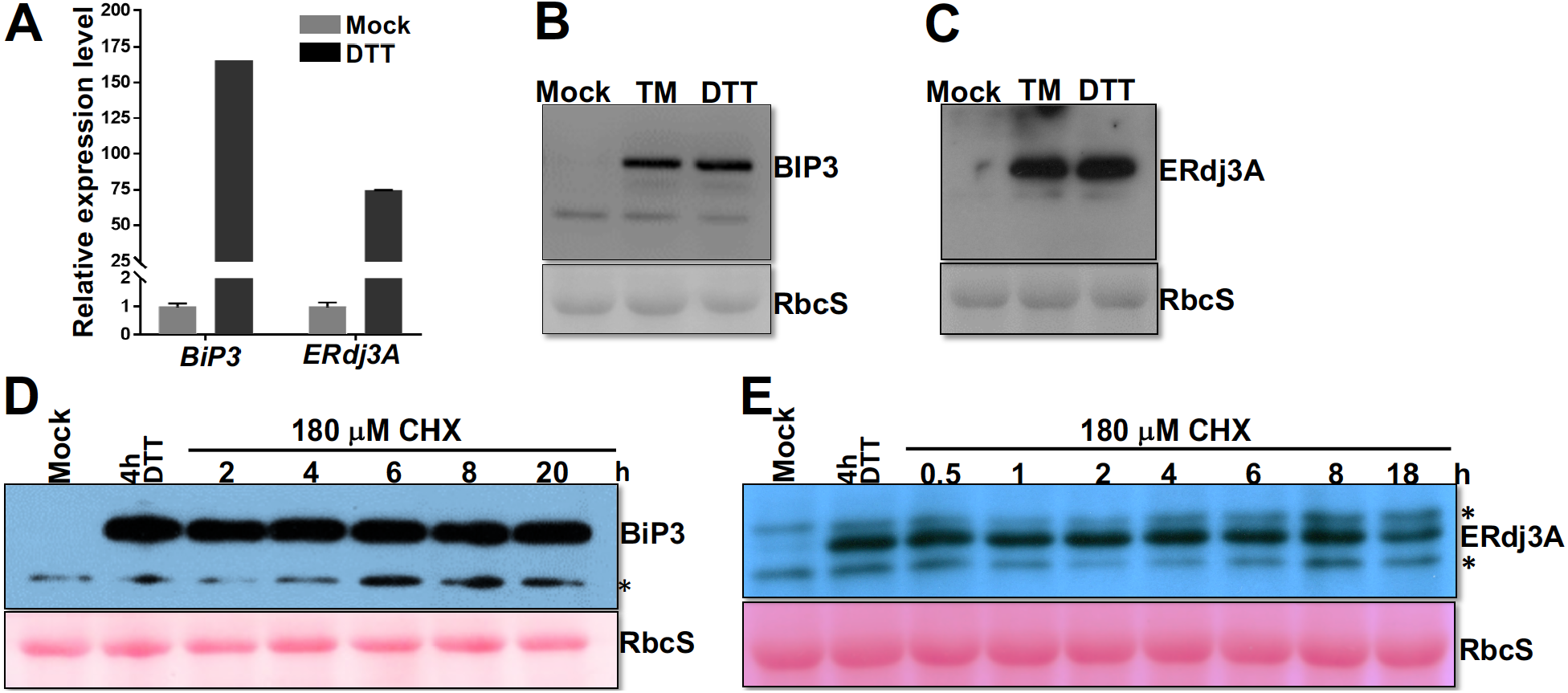
The UPR-induced BIP3 and TMS1/AtERdj3A are very stable proteins. **A**. A bar graph showing the transcript levels of *BIP3* and *ERdj3A* in DTT-treated seedlings in comparison to the mock-treated seedlings. **B-C.** Immunoblot analysis of the protein abundance of BIP3 (**B**) and ERdj3A (**C**) in wild-type seedings treated with or without TM or DTT. **E-F.** Immunoblot analysis of the protein abundance of BIP3 (**D**) and ERdj3A (**E**) in seedlings treated with DTT (for 4 hours) followed by 180 μM CHX (for varying durations). In **B**-**E**, the ponceau red-stained RbcS was used as the loading control.

## Discussion

In this study, we have shown that the endogenous TIN1 is an ER-localized protein that is rapidly degraded after its induction by UPR. Although an earlier study, which used protoplasts expressing a TIN1-GFP fusion protein, suggested that TIN1 was an ER-localized protein (Iwata et al., 2010), it remains unknown whether the endogenous TIN1 is also localized in the ER as the predicted TIN1 protein contains the N-terminal signal peptide but lacks a recognizable ER retention/retrieval motif. Besides, the C-terminal fusion of a GFP tag might prevent its packaging into trafficking vesicles for moving into the Golgi body. Because we have successfully generated a TIN1-specific antibody, we performed two biochemical assays to study the subcellular localization of the endogenous TIN1 protein in addition to confocal microscopic analysis of a TIN1-GFP fusion protein that was transiently expressed in tobacco leaf epidermal cells. The TIN1 protein was predicted to contain three N-glycosylation sites that were confirmed in this study by a simple Endo H assay of the endogenous TIN1 protein and analyses of Endo H and Kif sensitivity of 7 transgenically expressed TIN1-GFP fusion proteins lacking 1, 2, or all 3 predicted N-glycosylation sites. The simple Endo H assay was also used to confirm the ER localization of the endogenous TIN1 protein as its N-glycans could easily be deglycosylated by Endo H, which cleaves high mannose-type N-glycans of ER-localized or retained glycoproteins but not Golgi-processed complex-type N-glycans. The TIN1’s ER localization was further confirmed by the sucrose gradient ultracentrifugation in the presence or absence of Mg^2+^ that is required for the association of ribosomes with the ER (Lord et al., 1973). Given the lack of the HDEL ER retrieval motif, it is highly likely that TIN1 interacts with an ER-localized protein to stay in the folding compartment. Previous studies showed that EBS6/AtOS9, a well-established ERAD component, relies on its interaction with the ER membrane-anchored EBS5/AtHrd3 for its ER localization (Su et al., 2012). Similarly, SDF2 (stromal cell-derived factor-2) also forms a protein complex with AtERdj3B and BIP for its ER retention (Nekrasov et al., 2009). Identification of such TIN1-interacting proteins could help us to understand the likely biochemical function of TIN1 in coping with increased accumulation of misfolded proteins in the ER.

Our study demonstrated that TIN1 is an endogenous ERAD substrate that relies on its N-glycans and the Arabidopsis Hrd1-containing ERAD complex for its post-ER stress degradation. In addition, our experiments also suggested that TIN1 is rapidly degraded after its translation from stress-induced *TIN1* transcripts. We found that the heat treatment resulted in >30 fold induction of the *TIN1* transcript in vegetative tissues (**Supplemental Figure S1E**) but the TIN1 protein was below the detectable threshold by our anti-TIN1 antibody (**Supplemental Figure S13A**). By contrast, a similar heat treatment with the *hrd1a hrd1b* mutant seedlings or MG132-treated wild-type seedlings caused an easily observable accumulation of the TIN1 protein (**Supplemental Figure S13, A and B**). Previous studies suggested that two Arabidopsis proteins as potential endogenous ERAD substrates, AtOS9/EBS6 and UBC32 (Cui et al., 2012; Su et al., 2012; Chen et al., 2016; Chen et al., 2017). Both proteins are involved in ERAD with AtOS9/EBS6 functioning as a substrate recruitment factor that specifically recognizes a uniquely-demannosylated N-glycan on a terminally-misfolded glycoprotein (Huttner et al., 2012; Su et al., 2012) while UBC32 being an ER membrane-anchored E2 ubiquitin that works together with the ER membrane anchored E3 ligases (Cui et al., 2012). The discoveries of EBS6/AtOS9 and UBC32 being the endogenous ERAD substrate seemed to support the idea of the “ER tuning”, a term that was coined to describe the biochemical mechanism that regulates the abundance of the ERAD components to cope with dynamic changes of misfolded proteins in the ER in response to developmental cues and environmental signals (Bernasconi and Molinari, 2011). Our study revealed that ERAD is also used to degrade UPR-induced glycoproteins upon removal of ER stress. However, further study is needed to determine whether or not rapid degradation upon removal of ER stress is a unique property of TIN1 or applies to other known UPR-induced proteins. In mammalian cells, ERAD is known to regulate the abundance of two key sensors of the mammalian UPR mechanism, IRE1α and ATF6 (Horimoto et al., 2013; Sun et al., 2015), but little is known whether ERAD is also utilized to return the abundance of UPR-induced proteins to their pre-stress levels. Recent studies, however, suggested that a unique form of autophagy, known as “Recov-ER-phagy” is the most likely mechanism to not only reduce the abundance of UPR-induced ER proteins but also decrease the size of the ER that is expanded during the ER stress to increase ER folding capacity (Loi and Molinari, 2020). Identification of additional endogenous ERAD substrates will certainly shed more light on the physiological roles of the conserved degradation mechanism in plant growth and development.

What could be the signal that sends the endogenous TIN1 protein into the ERAD pathway? The 424-AA TIN1 polypeptide lacks any Cys residue and the DTT treatment unlikely affects its protein structure. However, it remains a possibility that the DTT treatment interferes with the TIN1 interaction with a yet to be discovered Cys-containing TIN1-binding protein, resulting in exposure of hydrophobic residues that could be detected by the Arabidopsis ERAD machinery. Our detected instability of TIN1-GFP in the *p35S::TIN1-GFP* transgenic line could be caused by an insufficient amount of such a TIN1-binding protein for the over-accumulated TIN1-GFP. Such a scenario is similar to what was observed previously for EBS6/AtOS9, which became unstable and was subject to ERAD-mediated degradation when its binding partner EBS5/AtHrd3A was mutated or become limited to the transgenically overproduced EBS6/AtOS9 (Su et al., 2012; Chen et al., 2017). EBS5/AtHrd3a not only works together with EBS6/AtOS9 to bring a terminally misfolded glycoprotein to the Hrd1-containing ERAD complex but is also necessary to retain EBS6/AtOS9, which lacks the HDEL ER-retrieval motif, inside the ER. By contrast, the endogenous EBS6/AtOS9 is a very stable protein that could still be detected 24 h after the wild type Arabidopsis seedlings were treated with CHX (**Supplemental Figure S16**). Thus, it is quite possible that the TIN1-binding protein that works together with TIN1 during the ER stress might be the same ones for binding and retaining TIN1 in the ER. Identification of TIN1-binding proteins should be one of the priorities for the future TIN1 study.

It is also important to note that both BIP3 and TMS1/AtERdj3A have 4 and 5 Cys residues, respectively, and are thus more prone to the DTT-triggered protein misfolding, yet both proteins remained stable after the DTT/CHX treatment. This revelation argued against the possibility of exposed hydrophobic residues being the main reason for driving the UPR-induced TIN1 but not UPR-induced BIP3 or TMS1/AtERdj3A in DTT-treated Arabidopsis seedlings into the ERAD pathway for rapid degradation. In mammalian cells, the degradation of a key-rate-limiting enzyme of the cholesterol biosynthesis pathway, HMGR (3-hydroxy-3-methyl-glutaryl-CoA reductase) that catalyzes reduction of HMG-CoA into mevalonate (Friesen and Rodwell, 2004), involves the binding of sterols to its membrane-embedded steroid-sensing domain and an adapter protein known as INSIG (insulin-induced gene) protein that interacts with ER membrane-anchored E3 ligases for HMGR ubiquitination (Johnson and DeBose-Boyd, 2018). It remains to be determined if TIN1 degradation involves a dedicated adapter protein or an ER stress-induced TIN1-binding chemical, which permits recognition of stress-induced TIN1 by the ERAD machinery in Arabidopsis. Alternatively, TIN1 might be an intrinsically disordered protein that dynamically changes its structure according to its binding partners but becomes an ERAD target when its binding partners disappear upon resolution of ER stress. Further investigation of rapid TIN1 degradation could uncover the mechanism(s) that drive some UPR-induced glycoproteins into the plant ERAD pathway and discover its binding partners important for its ER localization and its biochemical functions.

## Materials and Methods

### Plant materials and growth conditions

Most of the Arabidopsis wild-type, mutants, and transgenic lines are in the Columbia (Col-0) ecotype except for mutants and transgenic lines carrying the *bri1-5* mutation that was in the Wassilewskija-2 (Ws-2) ecotype. The Arabidopsis mutants used in this study include, *bri1-9* (Jin et al., 2007), *ebs5* (Su et al., 2011), *ebs6* (Su et al., 2012), *ebs7-1* (Liu et al., 2015)*, hrd1a hrd1b* (Su et al., 2011), *pawh1 pawh2* (Lin, et al 2019), *ire1a-2*, *ire1b-4* (Nagashima et al., 2011), *s2p*, *bzip28*, *bzip60* (Moreno et al., 2012). The T-DNA insertional mutants *CS411789* (*tin1-1*) and *CS808001* (*til1-1*) were obtained from Arabidopsis Biological Resource Center (ABRC) at Ohio State University. Seeds were surface sterilized using the ethanol-washing protocol and germinated seedlings were grown at 22°C in growth chamber or growth room under long-day (16h-light/8h-dark) photoperiodic condition.

### Generation of transgene constructs and transgenic plants

A 2-kb genomic DNA fragment upstream from the start codon of *TIN1* as the *TIN1* promoter was amplified from genomic DNAs of the wild-type Arabidopsis Col-0 seedlings using the *pTIN1-GUS* primer set (Table S1) and cloned into *pCambia1300* (https://cambia.org/welcome-tocambialabs/) to generate *pCambia1300-proTIN1::GUS* transgenes. The *35S::TIN1-GFP* and *35S::TIL1-GFP* transgene was created by cloning a 1,272-bp *TIN1* and 1,185-bp *TIL1* cDNA fragment amplified from the first-strand cDNAs converted from total RNAs of the wild-type Arabidopsis seedlings using the *cTIN1-GFP* and *cTIL1-GFP* primer sets (**Supplementary Table S1**) into the *pCambia1300p35S::C-GFP* vector, respectively. To generate the *pTIN1::TIN1-GFP*, a 3.3-kb *TIN1* genomic fragment containing the *TIN1* promoter was PCR amplified from the genomic DNAs of the wild-type Arabidopsis (Col-0) seedlings using the *gTIN1-GFP* primer set (**Supplementary Table S1**) and cloned into the *pCambia1300C-GFP* vector. The different N-glycan-mutant transgenes of *TIN1* was amplified from the *pTIN1::TIN1-GFP* plasmid using different site-directed mutagenesis primer sets (**Supplementary Table S1**). All created transgenes were fully sequenced to ensure no PCR-introduced sequence error, and were individually transformed into the *Agrobacterium tumefaciens* strain *GV3101* by electroporation and the resulting *Agrobacterial* strains were subsequently used to transformed into Arabidopsis plants using the floral-dipping method (Clough and Bent, 1998).

### Expression of fusion proteins and generation of antibodies

The first-strand cDNA preparation derived from total RNAs of wild-type Arabidopsis seedlings and the *TIN1-antigen* primer sets (**Supplementary Table S1**) were used to amplify a 1,191-bp *TIN1* cDNA fragment encoding a 396-AA (28-424) polypeptide. The amplified cDNA fragment was cloned into *pET-28a* (Novagen) and *pGEX-4T-1* (GE Healthcare) vectors, which were subsequently transformed into BL21-competent cells. The induction of His-and GST-fusion proteins and their subsequent purification using TALON^®^ Metal Affinity Resin (Clontech Laboratories) and Glutathione Sepharose^TM^ 4 Fast Flow beads (GE Healthcare), respectively, were carried out following the manufacturers’ recommended protocols. The purified His-TIN1 fusion protein was used to generate a custom anti-TIN1 antibody at MBL Beijing Biotech Co. (http://www.mbl-chinawide.cn), while the purified GST-TIN1 fusion protein was used to affinity-purify the custom-made anti-TIN1 antibody using an online protocol with nitrocellulose membrane (http://post.queensu.ca/~chinsang/lab-protocols/antibody-purification.html). The specificity of the purified anti-TIN1 antibodies was analyzed by immunoblotting with total proteins extracted from DTT treated 10-d-old seedlings of wild-type Arabidopsis plant and *tin1* mutants (**Supplementary Figure S2b**). The antibodies to BIP3 and TMS1 (ERdj3A) were custom-generated and affinity purified against the peptides, C^649^DPVIKSVYEKTEGENED^666^ and L^556^NGDIQFTKTRQKPQIK^572^, respectively, at ABclonal (www.abclonal.com.cn).

### Histochemical GUS staining assay

For histochemical GUS staining, 10-d-old *proTIN1::GUS* transgenic Arabidopsis seedlings were immersed into the GUS staining solution [0.1 M sodium phosphate buffer, 10 mM EDTA (pH 8.0), 0.5 mM K_3_Fe(CN)_6_, 0.5 mM K_4_Fe(CN)_6_, 1 mM X-glucuronide, 0.1% Triton X-100], vacuumed for 15 min, and subsequently incubated at 22°C with or without ER stress chemicals (5 μg/mL TM, 2 mM DTT, or 5 mM AZC) for overnight. GUS-stained tissues were dehydrated with 75% ethanol and observed by a ZEISS Stereo Discovery V8 microscope.

### Sucrose density-gradient centrifugation

Sixteen grams of 10-day-old Arabidopsis seedlings were ground in liquid N_2_ into a fine powder and immediately extracted by the homogenization buffer [50 mM Tris-HCl (pH 8.2), 20% (v/v) glycerol, 1 mM phenylmethylsulphonyl fluoride (PMSF, Sigma), 2 mM ethylenediaminetetraacetic acid (EDTA), 1 mM DTT, 2 protease inhibitor cocktail tablets (Roche) per 100 mL solution] at 4°C. The protein extracts were first filtered through Miracloth (CalBiochem) to remove insoluble plant debris and subsequently centrifuged at 5,000 x *g* for 5 min at 4°C to remove cellular debris and organelles. The supernatant was further centrifuged at 100,000 x *g* for 45 min to pellet the microsomes, which were resuspended in 1 mL resuspension buffer [25 mM Tris-HCl (pH 7.5), 10% (w/v) sucrose, 1 mM PMSF, 2 mM EDTA, 1 mM DTT, 2 protease inhibitor cocktail tablets (Roche) per 100 mL]. The microsomal resuspension was loaded onto the top of a 11 mL 20-50% (w/w) sucrose gradient in 10 mM Tris-HCl (pH 7.5), 2 mM EDTA, 1 mM DTT, 0.1 mM PMSF, and centrifuged at 100,000 x *g* for 16 h at 4°C. After centrifugation, 14 fractions (0.8 mL/each) were manually collected, and 50 µL protein sample for each fraction was mixed with 2x SDS buffer, 37°C warming for 1 hour, separated by 10% SDS/PAGE, and analyzed by immunoblotting. For the Mg^2+^-plus experiments, 5 mM MgCl_2_ was added to the buffers of homogenization, resuspension, and ultracentrifugation.

### Chemical treatment of Arabidopsis seedlings and subsequent immunoblot analyses

To analyze ER stress induction, 10-day-old Arabidopsis seedlings were carefully transferred into liquid ½ MS medium supplemented with 5 µg/mL tunicamycin (BioMol), 2 mM DTT (Sigma) and 5 mM AZC (TCI Shanghai), incubated for varying durations, and subsequently harvested into liquid nitrogen for immediate protein extraction or storage in a −80°C freezer. To study protein stability, 10-day-old Arabidopsis seedlings were pretreated with 2 mM DTT in liquid ½ MS medium for 4 hours and then carefully transferred into liquid ½ MS medium containing 180 µM CHX (Sigma), incubated for varying durations, and harvested into liquid nitrogen for protein extraction. To study the impact of Kif and MG132 treatment on protein stability, 10-day-old Arabidopsis seedlings were treated with 50 µM Kif (Abcam) or 80 µM MG132 (Sigma) in liquid ½ MS medium for varying durations and then harvested into liquid nitrogen for protein extraction. To study the impact of Endo H treatment on the electromobility of a given protein, 30 mg of 10-day-old Arabidopsis seedlings grown on ½ MS-agar medium were harvested into liquid nitrogen and ground thoroughly in 150 μL 2 x SDS loading buffer. An aliquot of 44 μL protein extracts was mixed with 1 μL Endo H-Hf and 5 μL 10x G5 buffer (New England Biolab), incubated at 37°C for 1 hour, and subsequently analyzed by SDS-PAGE and immunoblotting.

### Expression and confocal analysis of TIN1/TIL1 fusion proteins

Transgenes of *p35S::TIN1-GFP* and *p35S::TIL-GFP* were generated (see above) for analyzing the TIN1/TIL1 subcellular localization patterns. The plasmid *p35S::RFP-HDEL* was the same as previously described (Liu et al., 2015). To analyze protein interactions using the bimolecular fluorescence complementation (BiFC) assay, an 1,272-bp TIN1 cDNA amplified from the first-strand cDNAs of wild-type Arabidopsis seedlings using the *TIN1-NE* primer set (see **Supplementary Table S1**) was cloned into the *pVYNE* vector (Waadt et al., 2008). The same first-strand cDNA preparation was also used to amplify the full-length (1,872-bp) *MNS4* cDNA using the *MNS4-CE* primer set (**Supplementary Table S1**), which were subsequently cloned into the *pVYCE* vector (Waadt et al., 2008). These plasmids and the corresponding empty vectors, after verifying no PCR-introduced error by sequencing, were individually transformed into the *Agrobacterium tumefaciens* strain *GV3101*. The *p35S::RFP-HDEL*-carrying *GV3101* cells were mixed with the *p35S::TIN1-GFP* or *p35S::TIL1-GFP* -transformed *GV3101* strain and co-transformed into leaves of 3-week-old tobacco (*Nicotiana benthamiana*) plants by the agro-infiltration method (Yang et al., 2000). Similarly, a mixture of two *Agrobacterial* strains, one carrying a *pVYNE* plasmid and the other containing a *pVYCE* plasmid, was used to infiltrate young leaves of 3-week-old tobacco. Two days after infiltration, the transformed tobacco leaves were examined by confocal microscopy on a Leica SP8 (with LAS AF software, Leica Microsystems) for the subcellular localization patters of GFP-tagged TIN1/TIL1 and the ER-localized RFP-HDEL or the reconstituted YFP signal. GFP, YFP, and RFP were excited by using 488-, 514, and 542-nm laser light, respectively.

### RNA analyses

Ten-day-old Arabidopsis seedlings grown on ½ MS agar medium supplemented with or without certain chemicals were harvested and ground in liquid nitrogen into a fine powder and their total RNAs were extracted using the RNeasy Plant Mini Kit (QIAGEN). One microgram of the purified total RNAs were treated with RNase-free DNase I (TIANGEN) and subsequently reverse transcribed into the first-strand cDNAs by the iScript^TM^ cDNA synthesis Kit (Bio-Rad). The resulting cDNA preparations were used for traditional PCR or qRT-PCR analysis with gene specific oligonucleotides listed in **Supplementary Table S1**. The qPCR assays were performed on the CFX96 Real-Time System (Bio-Rad) with SYBR^®^ GREEN PCR Master Mix (Bio-Rad) following the manufacturer’s instruction. Three biological replicates with three technical repeats were conducted for each target mRNA with the *ACTIN8* cDNA being used as an internal reference to calculate the relative transcript abundance.

### Coimmunoprecipitation

Agro-infiltrated tobacco leaf disks were harvested and ground into fine powder in liquid N_2_. Total proteins were extracted into the immunoprecipitation (IP) buffer [50 mM Tris-HCl (pH 8.0), 150 mM NaCl, 5 mM EDTA, 0.1% (v/v) Triton X-100, 0.2% (v/v) Nonidet P-40 (Roche), 2 tablets of the protease inhibitor cocktail (Roche) per 100 mL], and centrifuged at 12,000 × *g* at 4 °C for 15 min. The supernatants were incubated with an anti-TIN1 antibody followed by protein-A/G-agarose beads (Abmart) or with anti-GFP Sepharose beads (Abcam) at 4 °C for 6 h. The resulting immunoprecipitants were subsequently washed five times with the IP buffer, resuspended in 2X SDS sample buffer, boiled at 95 °C for 10 min, separated by SDS/PAGE, and analyzed by immunoblotting.

### Scanning electron microscopic analysis of the pollen morphology

Arabidopsis pollen grains were mounted on stubs, coated with palladium-gold in a sputter coater (Emitech K575X, England), and then examined using a scanning electron microscope (Hitachi_S4700_FESEM, Japan) with an acceleration voltage of 5 kV.

## Acknowledgement

We thanked the ABRC for the seeds of the *tin1-1* (*CS411789*) and *til1-t* (*CS808001*) mutants and Jianxiang Liu (Zhejiang University) for the seeds of UPR mutants, and Shuh-ichi Nishikawa (Niigata University, Japan) for providing the anti-TMS1/AtERdj3A antibody. This work is partially supported by a grant from the National Natural Science Foundation of China (NSFC31730019) to JL and an annual research budget allocation of CAS to PW.

## Author Contributions

YW and JL designed experiments. YW, ZM, CZ, YC, LLin performed experiments and analyzed the data with technical and experimental support from JM, JZ, and LLiu. The first half of the project was supervised by LLiu while the second half was completed with supervision from PW. YW and JL wrote the manuscript with lots of inputs from JZ, LLiu, and PW.

## Data Availability

The data that supports the findings of this study are available from JL upon reasonable request.

## Supplementary Figures and Table

**Supplementary Figure S1. TIN1 is induced by ER stress and other stresses.**

**Supplementary Figure S2. The specificity test of a custom-made anti-TIN1 antibody.**

**Supplementary Figure S3. Neither *tin1-1* nor *TIN1*-overexpression had any observable effect on the pollen grain**

**Supplementary Figure S4. The presence of the TIN1-TIL1 pair in green organisms.**

**Supplementary Figure S5. TIL1 is also an ER-localized protein that does not respond to ER stress.**

**Supplementary Figure S6. Identification of a T-DNA insertional *til1-t* mutant.**

**Supplementary Figure S7. Overexpression of *TIN1* or *TIL1* had little impact on *bri1-9* mutant.**

**Supplementary Figure S8. The AZC-induced TIN1 protein is also rapidly degraded.**

**Supplementary Figure S9. TIL1 is a very stable protein.**

**Supplementary Figure S10. TIN1 can physically interact with MNS4 in plant cells.**

**Supplementary Figure S11. Conservation of the three glycosylated Asn residues between TIN1 and its homologs in green organisms.**

**Supplementary Figure S12. The effect of the known ERAD mutations on the *TIN1* transcript and TIN1 protein.**

**Supplementary Figure S13. The impact of heat stress, MG132, and *hrd1a hrd1b* double mutation on the TIN1 stability.**

**Supplementary Figure S14. The degradation of the AZC-induced TIN1 also involves the EBS7-PAWH1/2-Hrd1 machinery.**

**Supplementary Figure S15. *TIN1* is coexpressed with *BIP3* and *TMS1/ERdj3A*.**

**Supplementary Figure S16. The endogenous EBS6/AtOS9 is a very stable protein.**

**Supplementary Table S1. Oligonucleotide primers used in the study.**

